# Characterizing dysregulations via cell-cell communications in Alzheimer’s brains using single-cell transcriptomes

**DOI:** 10.1101/2023.07.16.548274

**Authors:** Che Yu Lee, Dylan Riffle, Yifeng Xiong, Nadia Momtaz, Ahyeon Hwang, Ziheng Duan, Jing Zhang

## Abstract

Alzheimer’s disease (AD) is a devastating neurodegenerative disorder affecting 44 million people worldwide, leading to cognitive decline, memory loss, and significant impairment in daily functioning. The recent single-cell sequencing technology has revolutionized genetic and genomic resolution by enabling scientists to explore the diversity of gene expression patterns at the finest resolution. Here, we leveraged the large-scale and publicly available single-nucleus RNA sequencing (snRNA-seq) in the human prefrontal cortex (PFC) from 23 AD samples and 13 controls to investigate cell-to-cell communication (C2C) in healthy brains and their perturbations in AD. Specifically, we first performed broad communication pattern analyses and discovered the inter-mixing of cell types and signaling pathways in AD brains. Secondly, we performed cell-type-centric analysis and found that excitatory neurons in AD have significantly increased their communications to inhibitory neurons, while inhibitory neurons and other supporting cells globally decreased theirs to all cells. Then, we delved deeper with a signaling-centric view, showing that canonical signaling pathways CSF, TGFβ, and CX3C are significantly dysregulated in their signaling to the cell type microglia/PVM and WNT pathway is dysregulated in its signaling from endothelial to neuronal cells in AD. Finally, after extracting 23 known AD risk genes, our intracellular communication analysis revealed a strong connection of extracellular ligand genes APP, APOE, and PSEN1 to intracellular AD risk genes TREM2, ABCA1, and APP in the communication from astrocytes and microglia to neurons. In summary, with the novel advances in single-cell sequencing technologies, we show that cellular signaling is regulated in a cell-type-specific manner and that improper regulation of extracellular signaling genes is linked to intracellular risk genes, connecting signaling to genetic differences manifested in AD.

**Author Summary:** Alzheimer’s is a devastating neurodegenerative disorder affecting 44 million people worldwide, leading to cognitive decline, memory loss, and significant impairment in daily functioning. The complex interplay of signaling genes suggests cells act in concert with other cells through cell-to-cell communication. Utilizing the recent advances in single-cell sequencing, we investigated dysregulated ligand-receptor gene pairs in the disease at the cell-type resolution. Specifically, our broad communication pattern analyses revealed the inter-mixing of cell types and signaling pathways in AD brains. Our cell-type-centric analysis found that excitatory neurons in AD have significantly increased their communications with inhibitory neurons, while inhibitory neurons and other supporting cells globally decreased theirs to all cells. With a signaling-centric view, we show that CSF, TGFβ, and CX3C pathways are significantly dysregulated in their signaling to the cell type microglia/PVM while the WNT pathway is dysregulated in its signaling from endothelial to neuronal cells in AD. Finally, our intracellular communication analysis revealed a strong connection of extracellular ligand genes APP, APOE, and PSEN1 to intracellular AD risk genes TREM2, ABCA1, and APP in the communication from astrocytes and microglia to neurons. In summary, performing a cell-to-cell communication analysis better explains the genetic differences manifested in Alzheimer’s.

## Introduction

Alzheimer’s disease is a devastating neurodegenerative disorder that affects 44 million people worldwide, leading to cognitive decline, memory loss, and significant impairment in daily functioning [1–3]. Understanding and studying Alzheimer’s disease is of utmost importance due to its widespread impact on individuals, families, and society as a whole [4–6]. Despite decades of efforts to narrow down several risk genes, the genetic and molecular mechanisms underlying AD are largely unknown. Bulk-tissue sequencing masks the heterogeneity of gene expression underlying distinct cell types [7]. As a result, we still face significant hurdles in developing effective treatment or prevention for this devastating disease.

The recent single-cell sequencing technology has revolutionized genetic and genomic studies by simultaneously profiling molecular signatures across thousands to millions of cells. It enables scientists to explore cellular diversity, gene expression patterns, and cellular interactions in complex tissues and health conditions, allowing us to identify unique cell types, discover disease-specific cellular signatures, and unravel the intricate mechanisms underlying genetic disorders [8–13]. As a result, several single-cell genomic research has been conducted to investigate the disease pathology and provides new molecular insights in AD research. For instance, Mathys et al. performed population-scale single-cell RNA sequencing (scRNA-seq) in post-mortem human brains from AD patients and healthy controls and revealed both cell-type-specific and cell-type-shared transcription perturbation signatures in AD [14]. On the other hand, Morabito et al. performed single-cell epigenetic and transcriptomic profiling and identified cell-type-specific cis-regulatory elements (CREs) and transcription factors (TF) that may mediate gene-regulatory changes in the late-stage AD [15]. Most of the existing studies solely focused on molecular perturbations within each individual cell. However, cells are not isolated entities but live in a microenvironment, or cell niche, composed of dynamically interacting entities, including extracellular matrix (ECM), neighboring cells, and soluble factors [8, 16, 17]. The complex and multidirectional interplay between these factors (and their properties) plays crucial roles in tissue development, cellular responses, disease progression, and therapeutic interventions [18–20]. Understanding and manipulating this relationship can provide insights into disease mechanisms and guide the development of novel therapeutic strategies.

To fill this gap, we leveraged the large-scale and publicly available single-nucleus RNA sequencing (snRNA-seq) in the human prefrontal cortex (PFC) to investigate cell-to-cell communication patterns in the human brain and their perturbations in AD patients. To achieve this goal, we first downloaded 36 single-cell nuclei RNA-seq (snRNA-seq) samples (23 AD and 13 control) and uniformly re-processed them with strict QCs. Next, we identified canonical cell types and their subclasses, consistent with cell types definitions from BICCN [21]. With such uniformly processed data, we built a high-confidence cell-to-cell communication network composed of signaling genes and inferred the major signaling pathway patterns in AD and healthy brains separately. Interestingly, we found that healthy brains form clear C2C patterns with distinct signaling usage, which has been significantly disrupted in AD brains. When compared to control, Alzheimer’s excitatory cell types seem to be sending more communication signals specifically to the inhibitory cell types, while inhibitory and supporting cell types globally decreased their outgoing signals to most cell types. We then delved deeper with a signaling-centric view and found that literature-canonical signaling pathways CSF, TGFβ, and CX3C are significantly dysregulated in their signaling to the cell type microglia/PVM while the WNT pathway is dysregulated in its signaling from endothelial to neuronal cells in AD [8, 22–30]. Finally, we calculated the regulatory scores of ligand genes and discovered, specifically, a strong connection of extracellular ligand genes APP, APOE, and PSEN1 to intracellular AD risk genes TREM2, ABCA1, and APP in the communication from astrocytes and microglia to neurons. In summary, with the novel advances in single-cell sequencing technologies, we show that cellular signaling is regulated in a cell-type-specific manner and that improper regulation of extracellular signaling genes is linked to intracellular risk genes, connecting signaling to genetic differences manifested in Alzheimer’s.

## Results

We utilized the data from the Accelerating Medicines Partnership Program for Alzheimer’s Disease Consortium (ADSP) [14]. We uniformly processed from the raw fastq files and kept 51,171 nuclei after strict QC (details in method section 4.1), including 31,294 nuclei from 23 AD samples and 19,877 nuclei from 13 healthy controls (**Fig. 1A**). These high-quality nuclei formed eight major cell types characterized by canonical marker genes (**Fig. 1B-C**). Since both excitatory and inhibitory neurons are composed of heterogeneous subclasses, we further clustered the neuronal clusters into subclasses by leveraging the existing BICCN reference dataset for PFC, including eight excitatory and nine inhibitory neuron subclasses (details in method section 4.2).

**Fig 1.**
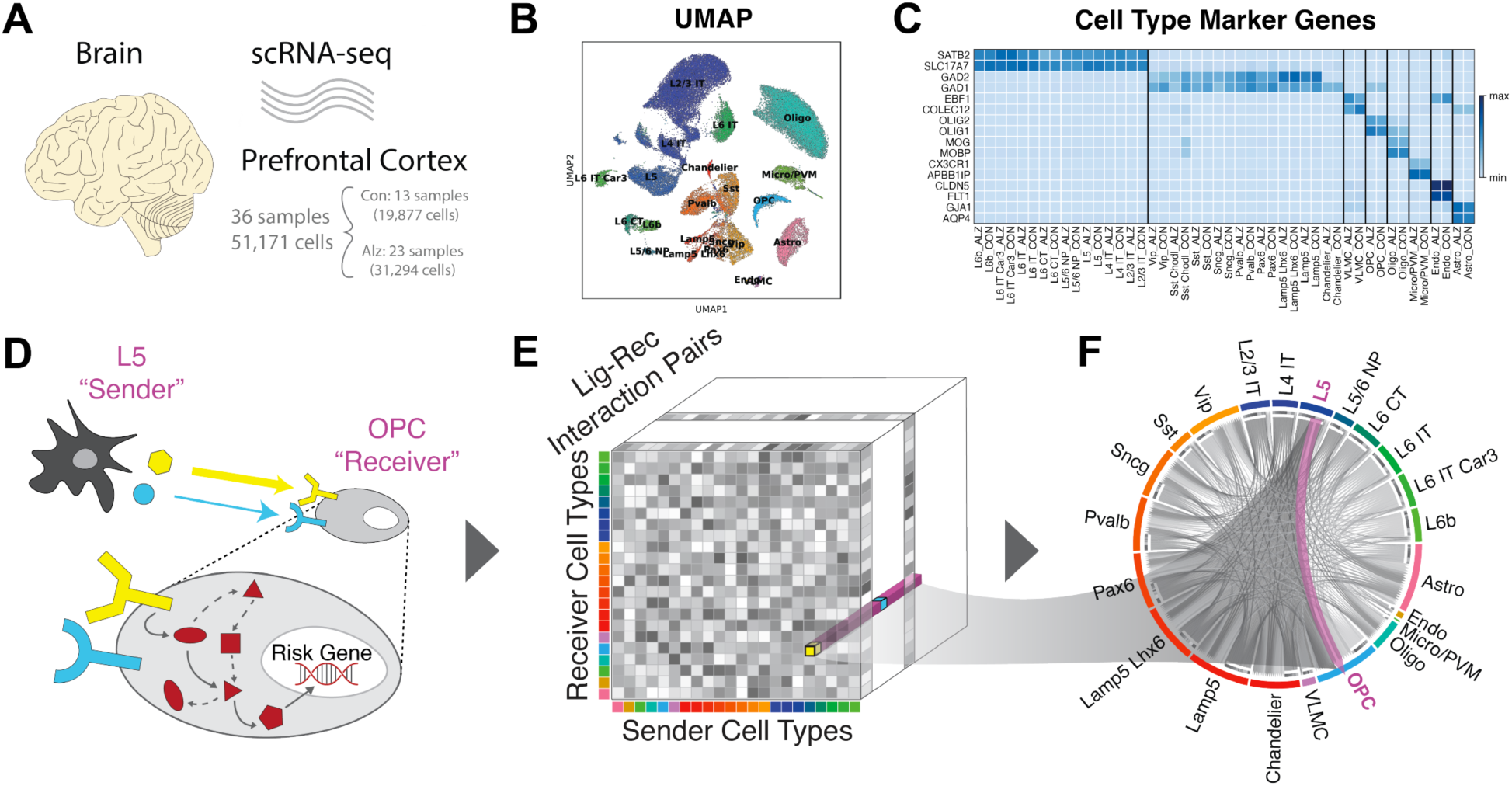
Data Overview. **(A)** Schematic of the Accelerating Medicines Partnership Program for Alzheimer’s Disease Consortium Data. **(B)** UMAP-embedding of the single-cell data labeled by cell type. **(C)** Table of marker genes for each cell type. **(D)** Schematic of the cell-to-cell communication analysis performed in the paper. **(E)** The data structure of our cell-to-cell communication analysis. It is a three-dimensional matrix representing the communication strength between any sender and receiver cell type pair via a specific ligand-receptor pair. **(F)** An overall cell-to-cell communication network that highlights the signaling from L5 neuron to OPC glial cell.

Next, we utilized the gene expression patterns of known ligand-receptor pairs from the snRNA-seq data to infer the C2C networks via a popular software package CellChat and connected them with downstream risk genes via NicheNet (**Fig. 1D**) [31, 32]. Specifically, we constructed a three-dimensional matrix representing the communication strength between any sender and receiver cell type pair via a specific ligand-receptor pair (**Fig. 1E**). As a result, this allowed us to aggregate the C2C communication patterns in AD and healthy controls, measure C2C changes between conditions, infer disease driving signal pathways, and connect risk genes to upstream ligand regulators in a cell-type-specific manner (**Fig. 1F**). We will discuss the detailed results in the following sections.

### 2.1 Communication Pattern Analysis reveals inter-mixing of cell types and signaling pathways in AD brains

With the 3D C2C matrix constructed, we first explored how multiple cell types coordinate intercellular communications using certain pathways in an unsupervised manner. To achieve this goal, we first flattened the 3D communication matrix into a 2D sender-by-LigandReceptorPair matrix and performed non-negative matrix factorization (NMF) to identify latent communication groups and their key ligand-receptor signaling contributors [31]. We demonstrated our outgoing C2C network results in the alluvial plot in **Fig. 2A**, where the middle bar represents the latent patterns, and the flow indicates how different signaling pathways (or cell types) belong to each pattern. Interestingly, we found normal brains employ three distinct outgoing communication latent patterns in three major cell groups, excitatory neurons, inhibitory neurons, and supporting cells (**left** panel in **Fig. 2A**). All of the outgoing supporting cells are characterized by pattern 1, dominated by biologically relevant pathways named after genes such as ANGPT, BMP, SPP1, and TGFβ [33–35] Inhibitory neurons are represented by pattern 2, driven by expected signaling pathways such as VIP, SST, CCK, and CRH [36–39] while excitatory neurons are characterized by pattern 3, driven by signaling pathways such as CSF, SEMA3, and NRG [40, 41]. These results show that biologically related cell types in normal brains rely on largely overlapping signaling networks. In contrast, we found that this pattern has been disrupted in AD brains (**right** panel in **Fig. 2A**). For instance, the inhibitory and excitatory neurons demonstrated mixed latent communication patterns (e.g., Chandelier cells have been grouped into excitatory patterns). In addition, the major driving signal pathways for different cell types also changed noticeably. For example, the WNT pathways became one major contributor to the excitatory group, while ANGPT switched from major contributors in supporting cells to the inhibitory group. Together, these results suggested extensive alterations in global C2C communication patterns and signaling usage in the outgoing network.

**Fig 2.**
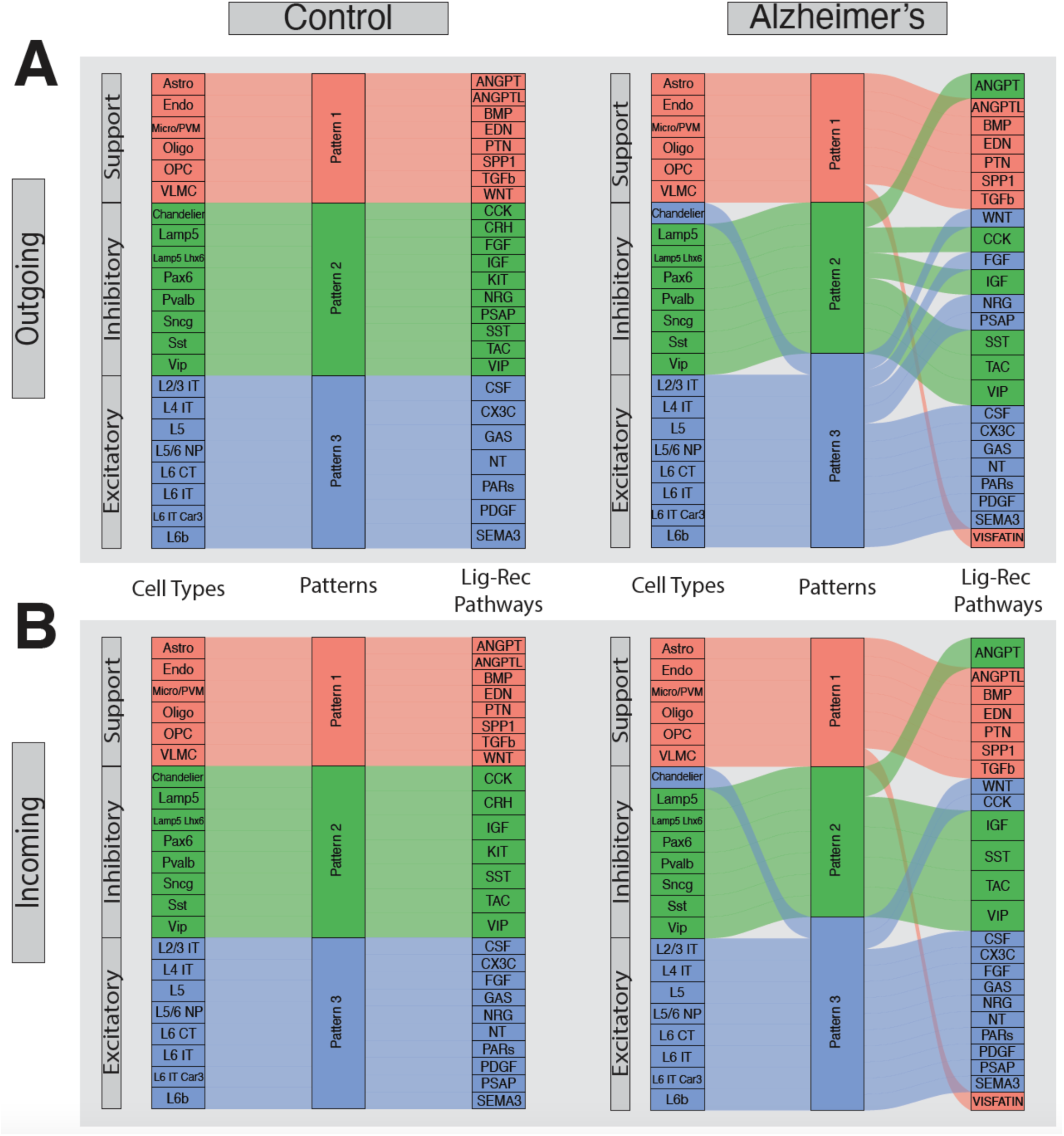
Pattern Analysis of Cell-to-Cell Communication. **(A)** Pattern analysis of the outgoing network for the control cells on the left and the Alzheimer’s cells on the right. The first column represents the cell types, while the last column represents the ligand-receptor pathways. Each pathway can contain multiple ligand-receptor pairs. **(B)** Pattern analysis of the incoming network for the control, healthy cells on the left and the Alzheimer’s cells on the right.

Next, we also explored the incoming C2C network latent patterns and their major signaling pathway contributors. Like the outgoing network, we found that normal brains demonstrate clear incoming latent communication patterns supported by biologically relevant signaling pathways (**left** panel in **Fig. 2B**), which has been disrupted in AD brains (**right** panel in **Fig. 2B**). For instance, we found that the inhibitory and excitatory groups have been mixed together (e.g., Chandelier cells). In general, the signaling pathways in both the outgoing and incoming networks are similar. Despite this, we do see small differences across the incoming and outgoing networks, specifically the FGF, NRG, and PSAP ligand-receptor interactions have switched from the inhibitory grouping to the excitatory one in the control samples, and the CCK and FGF signaling direction difference in the diseased ones (**Fig. 2A, B**). In summary, communication pattern analysis reveals the inter- mixing of cell types and signaling pathways in AD brains.

### 2.2 Pairwise cell type C2C comparison highlights disturbed communication strength across various cell types in AD brains

After checking the global C2C pattern perturbations, we focused on cell-type-centric communication changes by aggregating all Ligand receptor pairs in our 3D C2C matrix. First, we calculated the overall outgoing/incoming communication strength (*s*_*i*_) for a particular cell type *i* by aggregating all incoming/outgoing edges of each cell type and calculating the difference (Δ*s*_*i*_) between AD and control samples. Interestingly, we found that AD brains showed noticeably increased outgoing communication strength in most excitatory cell types (0.441 to 1.578, except for L2/3 IT), as compared to a general decrease in the inhibitory group (decrease of comm strength -0.148 to -1.383) (**Fig 3A1, S1 Table**). In the supporting cells, Astrocytes, OPC, and Micro/PVM also showed a higher level of outgoing communication strength (0.417, 0.933, 0.519, **Fig. 3A1**). Since each node in the circle plot represents a summation across all ligand-receptor pathways, we also indicated the boxplot of each individual ligand-receptor pathway for each cell type. Agreeing with the intensely bluely-colored “summed” node in its circle plot, the VLMC showed that more than 75% of its ligand-receptor pathways are down in Alzheimer’s compared to those of control (**Fig. 3A2**, **S2 Table**). On the other hand, in the incoming network, we found that the incoming communication strength is decreased in all excitatory cell types (-0.051 to -1.655), as compared to the general increase in the inhibitory group (an increase of comm strength 0.184 to 0.954), except for SST (**Fig. 3B1**, **S3 Table**). In the supporting cells, Astrocytes, Endo, and VLMC also showed a lower level of incoming communications (-0.338, -0.333, -0.605), except for a strong increase of Oligo (2.907, **Fig. 3B1**). Agreeing with the intensely redly-colored “summed” node in its circle plot, the Oligo showed that more than 75% of its ligand-receptor pathways are up in Alzheimer’s compared to those of control (**Fig. 3B2**, **S4 Table**).

**Fig 3.**
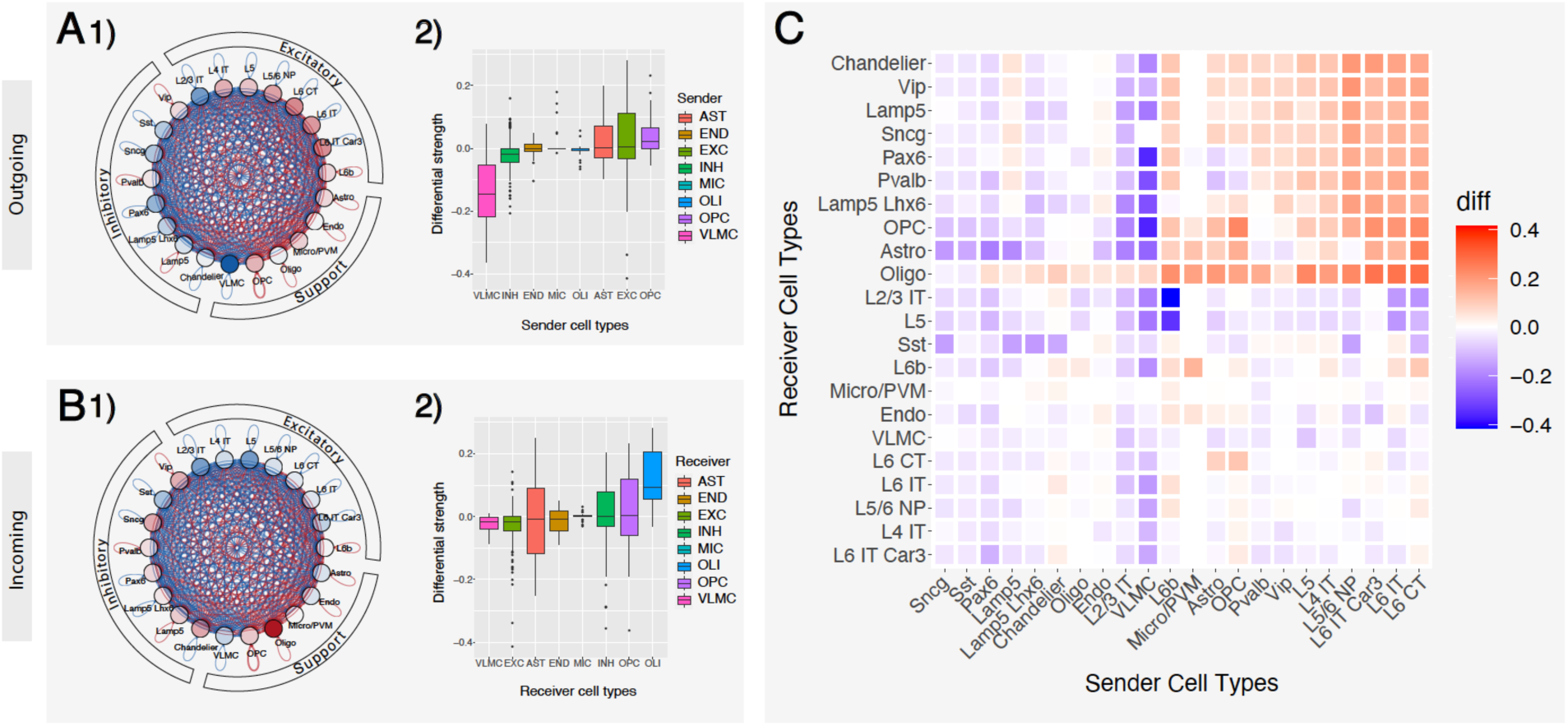
Cell-Type centric Cell-to-Cell Communication Analysis. **(A)** Differential network and boxplot of the outgoing cell-to-cell communication between Alzheimer’s and control. The nodes of the network were colored by the difference aggregated across signaling pathways. Red indicates an increased communication in Alzheimer’s (while blue indicates decreased). **(B)** Similar to A. Differential network and boxplot of the incoming cell-to-cell communication between Alzheimer’s and control. **(C)** Clustered heatmap of the cell-to-cell communication between all pair-wise cell types. Red indicates an increased communication in Alzheimer’s (while blue indicates decreased).

Next, we calculated the pair-wise AD-to-normal C2C communication, aiming to find the major driver cell types to explain the above changes. As shown in **Fig. 3C**, we found various cell types demonstrated distinct patterns of C2C disruption. Specifically, excitatory neurons demonstrated more targeted disruption in C2C communication patterns, while inhibitory neurons and supporting cells demonstrated more global disruptions. For example, the excitatory neuron groups significantly increased their communication mostly to inhibitory neurons (median Δ*s*_*i*_0.101, upper right red quadrant in **Fig. 3C**), but kept a similar level of communication to other cell types (median Δ*s*_*i*_ -0.006, bottom right quadrant in **Fig. 3C**). In contrast, most inhibitory neurons and supporting cells globally decreased their communication strengths to almost all cell types (median Δ*s*_*i*_ -0.034, left side of the heatmap in **Fig. 3C**). Our findings add to previous reports that AD patients showed an increased excitatory to inhibitory synaptic ratio by indicating direction in the communication [42, 43].

### 2.3 Canonical Neuroinflammation and Neuroprotection Signaling Pathways in AD are dis-regulated in a cell-type-specific manner

Our previous analyses mainly focused on the cell-type-level communication strength perturbations in the C2C network comparison without considering the impact of their communication pathways. To fill this gap, we also performed a signaling-pathway-centric analysis by evaluating the contribution of all involved ligand-receptor pairs (details in Method section 4.3). Each signaling pathway contains multiple ligand-receptor gene pairs. As shown in **Fig. 4A**, many robustly expressed pathways in the human brain demonstrated significantly altered involvement in C2C communication network. For example, for each pathway, we aggregated the communication strength of all involved Ligand Receptor pairs and across all cell type pairs. After permutation tests, we found that 22 pathways showed significantly decreased communication activity, while 2 pathways demonstrated increased activity (**Fig. 4A**). We chose to focus on 4 canonical, literature-driven ligand-receptor interactions for further analysis, namely the WNT, CSF, TGFβ, and CX3C pathways (**Fig. 4B-E**).

**Fig 4.**
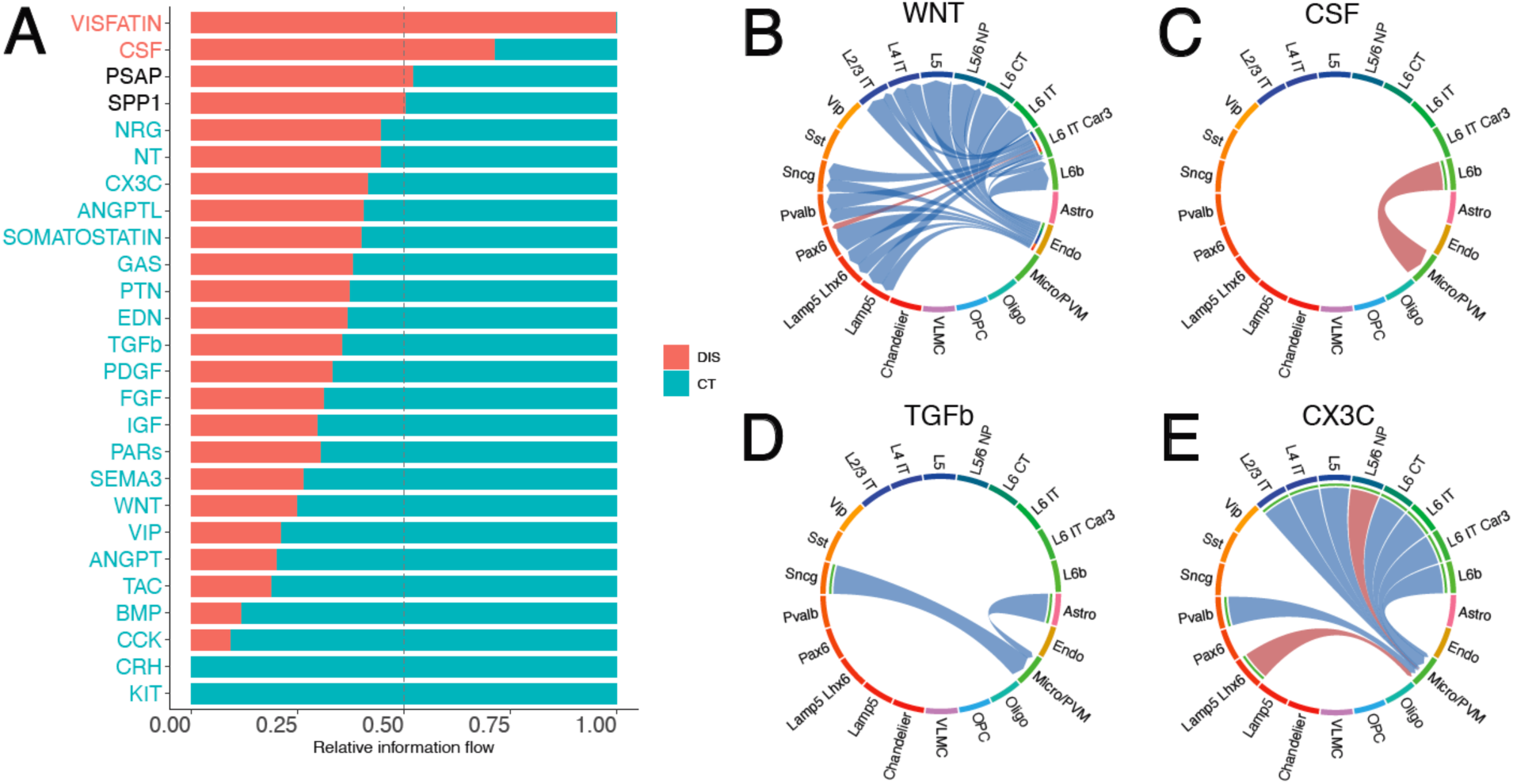
Signaling Pathways of Cell-to-Cell Communication Analysis. **(A)** Comparison of the signaling pathway flow between Alzheimer’s and control. The flow is defined as the summation of the communication strength of that specific pathway across all sender-receiver cell type pairs. Red indicates an increased communication in Alzheimer’s (while blue indicates decreased). **(B)** Communication strength difference among cell types in the pathway WNT. Red indicates an increased communication in Alzheimer’s (while blue indicates decreased). **(C)** Communication strength difference among cell types in the pathway CSF. **(D)** Communication strength difference among cell types in the pathway TGFβ. **(E)** Communication strength difference among cell types in the pathway CX3C.

Neuronal inflammation plays a significant role in the AD pathology. For instance, immune cells such as microglia respond to the accumulation of beta-amyloid plaques, a hallmark of Alzheimer’s, by triggering an inflammatory response [44]. Also. prolonged microglia activation can result in chronic inflammation, leading to neuronal damage and the exacerbation of plaque buildup, thus creating a vicious cycle [45]. Consistently, we found that two inflammation-related pathways WNT and CSF are dysregulated in the C2C communication process in our analysis. For example, the WNT signaling pathway plays multifaceted roles in CNS diseases by modulating neuroimmune interactions [46]. We found that the WNT pathway has significantly reduced its involvement in C2C communication (30% of control, P=2.086e-7, **Fig. 4A**, **S5 Table**), which has been primarily driven by the global reduction of communication usage from the sender endothelial cells to both inhibitory and excitatory neuron receivers (**Fig. 4B**). Mechanistically, the downregulation of the WNT ligand gene may cause overactivity of the lithium-targeted GSK3β enzyme, leading to changes in neurogenesis, inflammation, oxidative stress, and circadian dysregulation in neuronal cell types [24]. Additionally, lines of literature reported that the CSF pathway is a well-known disease-related signaling pathway primarily involved in microglia [22, 36, 47, 48]. which can activate the recruitment of microglia and worsen inflammatory response. Consistently, we found that the CSF pathway has been significantly upregulated in AD patients (2.5x of control, P=0, **Fig. 4A**). Such increased involvement is mainly driven by the increased communication from the excitatory neurons L6b to Microglia cells (**Fig. 4C**).

Next, we move on to neuroprotective signaling pathways. We observed the downregulation of TGFβ signaling in Alzheimer’s in the communication to Micro/PVM cell type (60% of control, P=0, **Fig. 4A, D**, **S5 Table**). A decreased TGFβ1 has been associated with a higher burden of Aβ in the parenchyma, which correlates with an increased microglia activation [49]. The suppression of the neuroprotective role of the signaling pathway TGFβ1 against Aβ toxicity in the diseased cell types may be the molecular mechanism underlying the symptoms of Alzheimer’s disease. Adding on, we also found the decrease of another neuroprotective signaling pathway, the CX3C pathway (70% of control, P=4.883e-2, **Fig. 4A**). CX3CL1 has been demonstrated to play a neuroprotective role in CNS by reducing neurotoxicity and microglial activation [50]. Our C2C analysis agrees with the literature as we see all communication in the CX3C pathway is directed to the Micro/PVM cell type. Moreover, with single-cell resolution, we are able to further see that this decrease happens primarily from the excitatory neurons to Micro/PVM (**Fig. 4E**). In summary, we discover that both the signaling pathways that cause neuroinflammation and those that protect against it are regulated in a cell-type-specific manner.

**Fig 5.**
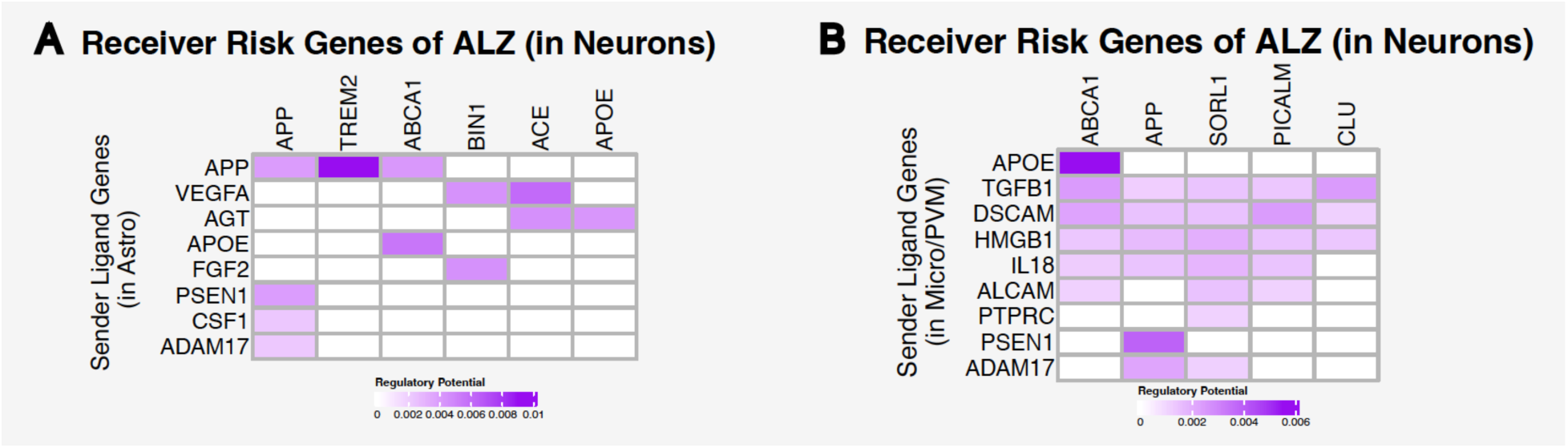
Connection to Intracellular Risk Genes. **(A)** Connection of astrocyte’s ligand genes to neurons’ risk genes in Alzheimer’s. More intensely colored boxes indicate a stronger regulation. **(B)** Connection of microglia/perivascular macrophages’ ligand genes to neurons’ risk genes in Alzheimer’s.

### 2.4 Intracellular Cell-to-Cell Communication Analysis reveals a strong connection to Neuroinflammatory AD risk genes

Next, we seek to see how that extracellular signaling is precisely connected with known AD risk genes in an intra-cellular manner. To accomplish this, we first extracted AD risk genes, including APP, ABCA1, and TREM2 (details see method section 4.4). Then, we defined Ligand-to-risk-gene regulation scores by combining C2C communication networks across cells and gene-gene interaction networks within cells via NicheNet [51]. Since many of the well-known disease risk genes appear to be regulated in a cell type-specific fashion by our extracellular cell-to-cell communication analysis above, we considered only a subset of sending cell types and receiving cell types (**Fig. 5**). Specifically, we set astrocytes and microglia cells as the sender cell types and neurons as the receiver cell types to find neuroinflammatory signals that cause neuron dysfunctions or neuronal death.

In the astrocyte-to-neuron signaling, we find ligand-target links connecting neurological risk genes to potential upstream effectors, such as the APP-TREM2 (Regulatory Score 0.00904, the maximum) and APP-ABCA1 link (Regulatory Score 0.00410) (**Fig. 5A**). TREM2 in microglia and macrophages results in decreased phagocytosis of apoptotic neurons, increasing Aβ accumulation in AD Phenotype [36, 52, 53]. Additionally, ABCA1 deficiency increased amyloid deposition in the brains of amyloid precursor protein (APP) transgenic mice [54, 55]. In the microglia-to-neuron signaling, we find ligand-target links such as APOE-ABCA1 (Regulatory Score 0.005490, the maximum), again, and PSEN1-APP (Regulatory Score 0.003858) (**Fig. 5B**). PSEN1 and PSEN2 mutations have been linked with the Amyloid protein precursor in early-onset Alzheimer disease [56]. By linking our extracellular signaling above to intracellular risk genes, our ligand-target analysis completes our cell-to-cell communication analysis and supports our hypothesis that AD communication dysregulations happen with cell-type-specificity and that improper regulation of extracellular signaling genes is linked to intracellular risk genes.

## Discussion

Alzheimer’s Disease is a neurological disorder involving genetic, epigenetic, and environmental factors through various processes. The improper regulation of cell-cell signaling can be linked to Alzheimer’s disease [57–59]. With the explosion of data in the recent consortium initiatives and the recent technological developments in the single-cell revolution of investigating AD pathology at the finest possible resolution – individual cells, we are able to, for the first time, systematically compare cell-cell communications in Alzheimer’s and control brains.

Results from our C2C analysis have shown that there is a global C2C communication pattern intermixing (inhibitory and excitatory groups **Fig. 2**) and signaling pathway misusage in AD brains (e.g. ANGPT, WNT, CCK pathways, **Fig. 2**). Additionally, we also observed a large degree of C2C communication disruption heterogeneity across various cell types. For example, excitatory neurons tend to solely increase their communication strength with inhibitory neurons, while supporting cells and inhibitory neurons globally decrease their communication to most cell types (**Fig. 3**). This signifies the importance of employing single cell technologies in AD study to dissect the extensive genetic heterogeneity in complex tissues like the human brains. Furthermore, we highlighted the involvement of the neural inflammatory and neural protective pathways, such as WNT, CSF, TGFβ, and CX3C, in AD patients from the C2C communication aspect (**Fig. 4**). Their disturbed behavior can pass erroneous information both inter- and intra cells to directly impact well-known AD risk genes.

In this study, cell-cell communication is inferred from the expression of protein-coding genes, thus not fully capturing other signaling events in the brain, such as nonprotein molecules (e.g., neurotransmitters). We seek to address this limitation in the future by considering also the gene expression of the synthesis and transporter proteins for neurotransmitters to study more thoroughly the communication networks of the brain. Another limitation of the study, as in other cell-cell interaction studies based on scRNA-seq or snRNA-seq, lies in the lack of spatial information [60]. Finally, in our last risk gene analysis, we can provide a finer resolution of our single cells by also incorporating the chromatin-accessibility analysis scATAC-seq [61].

Despite these limitations, we believe our work can serve as a valuable first step in investigating the intercellular molecular mechanisms underlying Alzheimer’s disease beyond the general bulk-seq picture. With further technological advances and community efforts for population-scale single-cell sequencing, we expect exponentially increased power to accurately quantify C2C communication and its alterations in AD brains, hoping for the subsequent alleviation of the pain caused by AD.

## Supporting information captions

**S1 Table. Aggregated Communication for the Outgoing Network.**

**S2 Table. Distribution of Communication for the Outgoing Network. S3 Table. Aggregated Communication for the Incoming Network.**

**S4 Table. Distribution of Communication for the Incoming Network. S5 Table. Ratio and Pvalues of each Communication Pathway.**

**S6 Table. Alzheimer’s Risk Genes.**

