## Supplementary Tables for "Characterizing dysregulations via cell-cell communications in Alzheimer’s brains using single-cell transcriptomes"

### S1 Table. Aggregated Communication for the Outgoing Network.

| | |
|---|---|
| **L6b** | 0.441297442205382 |
| **L6 IT Car3** | 1.49872281745899 |
| **L6 IT** | 1.32054270489425 |
| **L6 CT** | 1.57856747984515 |
| **L5/6 NP** | 1.19518432471581 |
| **L5** | 0.693334680836904 |
| **L4 IT** | 0.973861153931422 |
| **L2/3 IT** | -1.92656006857613 |
| **Vip** | 0.376883866018848 |
| **Sst** | -0.912839763723862 |
| **Sncg** | -1.00091224331024 |
| **Pvalb** | 0.237634791621928 |
| **Pax6** | -1.38360063501575 |
| **Lamp5 Lhx6** | -0.804932299022804 |
| **Lamp5** | -0.433006624514807 |
| **Chandelier** | -0.14814122505262 |
| **VLMC** | -3.36286670165078 |
| **OPC** | 0.933437845741916 |
| **Oligo** | -0.143111443345204 |
| **Micro/PVM** | 0.518772701803891 |
| **Endo** | -0.0696092874028197 |
| **Astro** | 0.417340482540535 |

### S2 Table. Distribution of Communication for the Outgoing Network.

| name | 10 | 25 | 50 | 75 | 90 |
|---|---|---|---|---|---|
| AST | -0.057893006 | -0.029932685 | 0.001072532 | 0.070534979 | 0.089582613 |
| END | -0.039509167 | -0.008774243 | 0 | 0.01436848 | 0.027855213 |
| MIC | 0 | 0 | 0 | 0 | 0.101449972 |
| OPC | -0.030851994 | 0 | 0.023003375 | 0.064671333 | 0.144084087 |
| OLI | -0.044075282 | -0.00829059 | 0 | 0 | 0.005027021 |
| VLMC | -0.288846693 | -0.216892145 | -0.144233731 | -0.051729043 | 0 |

### S3 Table. Aggregated Communication for the Incoming Network.

| | |
|---|---|
| **L6b** | -0.0507919047368934 |
| **L6 IT Car3** | -0.657613323881614 |
| **L6 IT** | -0.294521119254933 |
| **L6 CT** | -0.23412574709297 |
| **L5/6 NP** | -0.35579860223482 |
| **L5** | -1.65548802920828 |
| **L4 IT** | -0.447851755989095 |
| **L2/3 IT** | -1.59418302409378 |
| **Vip** | 0.85997846333682 |
| **Sst** | -0.957502365114291 |
| **Sncg** | 0.905339181270896 |
| **Pvalb** | 0.301809851791979 |
| **Pax6** | 0.362277369514244 |
| **Lamp5 Lhx6** | 0.183606443437312 |
| **Lamp5** | 0.496960796762695 |
| **Chandelier** | 0.953683548614192 |
| **VLMC** | -0.605220395717914 |
| **OPC** | 0.549449701930449 |
| **Oligo** | 2.9066157679128 |
| **Micro/PVM** | 0.00377748898219819 |
| **Endo** | -0.332622021153463 |
| **Astro** | -0.337780325075515 |

### S4 Table. Distribution of Communication for the Incoming Network.

| name | 10 | 25 | 50 | 75 | 90 |
|---|---|---|---|---|---|
| AST | -0.057893006 | -0.029932685 | 0.001072532 | 0.070534979 | 0.089582613 |
| END | -0.039509167 | -0.008774243 | 0 | 0.01436848 | 0.027855213 |
| MIC | 0 | 0 | 0 | 0 | 0.101449972 |
| OPC | -0.030851994 | 0 | 0.023003375 | 0.064671333 | 0.144084087 |
| OLI | -0.044075282 | -0.00829059 | 0 | 0 | 0.005027021 |
| VLMC | -0.288846693 | -0.216892145 | -0.144233731 | -0.051729043 | 0 |

### S5 Table. Ratio and Pvalues of each Communication Pathway.

| | name | contribution | contribution.scaled | group | contribution.relative.1 | pvalues |
|---|---|---|---|---|---|---|
| **VISFATIN1** | VISFATIN | 0.633692405253888 | 2.19206136978575 | DIS | Inf | 0 |
| **CSF1** | CSF | 0.03198058526324 | 0.290475910345929 | DIS | 2.5 | 0 |
| **PSAP1** | PSAP | 6.7190932181216 | 18.2294855930064 | DIS | 1.1 | 0.898337619940568 |
| **SPP11** | SPP1 | 0.429365695226487 | 1.18280725301718 | DIS | 1 | 0.84375 |
| **NT1** | NT | 0.829284123886454 | 5.3420957464412 | DIS | 0.8 | 0.000164031982421875 |
| **NRG1** | NRG | 111.389458388842 | 18.8691166664452 | DIS | 0.8 | 9.543754933973E-36 |
| **CX3C1** | CX3C | 0.144752418820109 | 0.517402723586939 | DIS | 0.7 | 0.048828125 |
| **SOMATOSTATIN1** | SOMATOSTATIN | 0.142258582185443 | 0.512791881654416 | DIS | 0.7 | 0.0078125 |
| **ANGPTL1** | ANGPTL | 0.37661846358958 | 1.02404180119196 | DIS | 0.7 | 0.00558090209960938 |
| **EDN1** | EDN | 0.00703327838561526 | 0.201730755485365 | DIS | 0.6 | 0 |
| **TGFb1** | TGFb | 0.0194877700632485 | 0.253938059593817 | DIS | 0.6 | 0 |
| **GAS1** | GAS | 0.920062925258781 | 12.0028978465287 | DIS | 0.6 | 4.50204691572963E-06 |
| **PTN1** | PTN | 6.08238335869453 | 17.5898545195675 | DIS | 0.6 | 1.17730890161945E-11 |
| **PDGF1** | PDGF | 0.587378213327422 | 1.87939419659463 | DIS | 0.5 | 1.81898940354586E-12 |
| **FGF1** | FGF | 1.64159451337309 | 15.0313302258123 | DIS | 0.5 | 4.83815056163615E-18 |
| **PARs1** | PARs | 0.924807139583553 | 12.7926214687764 | DIS | 0.4 | 7.10542735760102E-15 |
| **IGF1** | IGF | 2.56571978965721 | 15.9907768359705 | DIS | 0.4 | 7.22758378973869E-26 |
| **SEMA31** | SEMA3 | 2.21057117529351 | 15.6709612992511 | DIS | 0.4 | 3.37240084539392E-14 |
| **ANGPT1** | ANGPT | 0.00968541778505352 | 0.215650449883497 | DIS | 0.3 | 0 |
| **VIP1** | VIP | 0.055800712620159 | 0.346504111856335 | DIS | 0.3 | 0.03125 |
| **WNT1** | WNT | 0.211700658453888 | 0.644088367699347 | DIS | 0.3 | 2.08616256713867E-07 |
| **TAC1** | TAC | 0.0540843906907821 | 0.342793325113808 | DIS | 0.2 | 0.0390625 |
| **CCK1** | CCK | 0.076322726606188 | 0.388683929686567 | DIS | 0.1 | 1.86264514923096E-09 |
| **BMP1** | BMP | 0.400871712855227 | 1.09395567281311 | DIS | 0.1 | 3.88933214472469E-15 |
| **CRH1** | CRH | 0 | 0 | DIS | 0 | 0 |
| **KIT1** | KIT | 0 | 0 | DIS | 0 | 0 |

### S6 Table. Alzheimer's Risk Genes.

| |
|---|
| ABCA7 |
| EIF2AK2 |
| APP |
| CR1 |
| FERMT2 |
| APOE |
| CLU |
| PICALM |
| ADAM10 |
| ABCA1 |
| HLA-DRB1 |
| CD33 |
| CD2AP |
| MME |
| TREM2 |
| CASS4 |
| MS4A4A |
| APH1B |
| BIN1 |
| ACE |
| SORL1 |
| GRN |
| SNX1 |
